# The Metaxin Mitochondrial Import Proteins: Multiple Metaxin-like Proteins in Fungi

**DOI:** 10.1101/2022.01.04.474931

**Authors:** Kenneth W. Adolph

## Abstract

Multiple metaxin-like proteins are shown to exist in fungi, as also found for the metaxin proteins of vertebrates and invertebrates. In vertebrates, metaxins 1 and 2 are mitochondrial membrane proteins that function in the import of proteins into mitochondria. Fungal metaxin-like proteins were identified by criteria including their homology with human metaxins and the presence of characteristic GST_N_Metaxin, GST_C_Metaxin, and Tom37 protein domains. Fungi in different taxonomic divisions (phyla) were found to possess multiple metaxin-like proteins. These include the Ascomycota, Basidiomycota, Blastocladiomycota, Chytridiomycota, Mucoromycota, Neocallimastigomycota, and Zoopagomycota divisions. Most fungi with multiple metaxin-like proteins contain two proteins, designated MTXa and MTXb. Amino acid sequence alignments show a high degree of homology among MTXa proteins, with over 60% amino acid identities, and also among MTXb proteins of fungi in the same division. But very little homology is observed in aligning MTXa with MTXb proteins of the same or different fungi. Both the MTXa proteins and MTXb proteins have the protein domains that characterize the metaxins and metaxin-like proteins: GST_N_Metaxin, GST_C_Metaxin, and Tom37. The metaxins and metaxin-like proteins of vertebrates, invertebrates, plants, protists, and bacteria all possess these domains. The secondary structures of MTXa and MTXb proteins are both dominated by similar patterns of α-helical segments, but extensive β-strand segments are absent. Nine highly conserved α-helical segments are present, the same as other metaxins and metaxin-like proteins. Phylogenetic analysis reveals that MTXa and MTXb proteins of fungi form two separate and distinct groups. These groups are also separate from the groups of vertebrate metaxins, metaxin-related Sam37 proteins of yeasts, and metaxin-like FAXC proteins.

## 1. INTRODUCTION

The metaxins were originally identified in human and mouse as mitochondrial membrane proteins that function in protein import into mitochondria. In particular, β-barrel membrane proteins in the cytoplasm are taken up and integrated into the mitochondrial outer membrane. Two vertebrate metaxins were initially identified – metaxins 1 and 2 – and a third metaxin has been found to be widely distributed in vertebrates (Adolph, 2019). Proteins homologous to vertebrate metaxins 1 and 2, but not metaxin 3, have also been identified in invertebrates (Adolph, 2020a). Among many examples of invertebrates with metaxin 1 and 2 proteins are the nematode *C. elegans*, a model organism of biology research, and the sea urchin *Strongylocentrotus purpuratus*, also a model research organism. In addition, insects have proteins homologous to vertebrate metaxins 1 and 2. Examples are the fruit fly *Drosophila melanogaster*, widely studied in genetics research, and *Apis mellifera*, the most common domesticated honey bee.

Plants and bacteria were found to possess proteins that are metaxin-like but not directly homologous to vertebrate metaxin 1, 2, or 3 (Adolph, 2020b). A wide variety of plant species contain metaxin-like proteins, including plants of economic importance such as *Zea mays* (maize or corn) and *Glycine max* (soybean). Metaxin-like proteins are also present in bacteria, mainly in bacteria of the Proteobacteria phylum. As examples, metaxin-like proteins are found in *Francisella tularensis*, a human pathogen, and *Colwellia psychrerythraea*, an obligate psychrophile (thrives in permanently cold temperatures). Protists and fungi likewise possess metaxin-like proteins (Adolph, 2021). Protists with these proteins include animal-like protists such as protozoa and amoebae, and plant-like protists such as phytoplankton and algae. Fungal metaxin-like proteins exist in species representing the major taxonomic divisions of Ascomycota, Basidiomycota, Blastocladiomycota, Chytridiomycota, Mucoromycota, Neocallimastigomycota, and Zoopagomycota.

For fungi, as for other organisms, genome sequencing has provided profound insight into the evolution and functioning of the species. It was therefore primarily fungi with fully sequenced genomes that were selected for this study. Model systems used in biological research also received high priority, as did organisms of economic or medical importance. An example is *Schizosaccharomyces pombe* fission yeast, a member of the Ascomycota division. *S. pombe* is a model organism of molecular and cell biology research, and was among the first organisms to have its genome sequenced (Wood et al., 2002). Another important research organism of the Ascomycota division is *Aspergillus nidulans*, also with a sequenced genome (Galagan et al., 2005). The ascomycete fungus *Penicillium rubens* (genome sequence: van den Berg et al., 2008) is used in the industrial production of penicillin. *Botrytis cinerea* is an ascomycete fungus of economic significance as a wine grape fungus that produces “gray rot” and “noble rot” (sequence: Amselem et al., 2011). Other ascomycete fungi, some identified as human pathogens, were also included in this study: *Blastomyces dermatitidis* (Munoz et al., 2015), *Histoplasma capsulatum* (Sharpton et al., 2009), *Microsporum canis* (Martinez et al., 2012), *Paecilomyces variotii* (Urquart et al., 2018), and *Talaromyces marneffei* (Nierman et al., 2015).

The aim of this study was to identify fungi with multiple metaxin-like proteins, as found for the metaxin proteins of vertebrates and invertebrates. Fungi of the Ascomycota division, mentioned above, were the first revealed to possess multiple metaxin-like proteins, designated MTXa and MTXb. In addition to fungi of the Ascomycota division, fungi of other major divisions were also found to possess multiple metaxin-like proteins. The fungi chosen for this study are representative of fungi in those divisions. Specifically, the Basidiomycota division is represented by, among other fungi, *Neolentinus lepideus* (Nagy et al., 2016), which is a mushroom that grows on decaying wood. A Blastocladiomycota representative is *Catenaria anguillulae* (Mondo et al., 2017), which is a parasite of nematodes that are parasites of crop plants. And Chytridiomycota fungi include *Gonapodya prolifera* (Chang et al., 2015) and *Rhizoclosmatium globosum* (Mondo et al., 2017). *Glomus cerebriforme* (Morin et al., 2019), which establishes a symbiotic relationship with plant roots (mycorrhizas), represents the Mucoromycota division. *Neocallimastix californiae* and *Piromyces finnis* (Haitjema et al., 2017), found in the digestive tract of large herbivores, represent the Neocallimastigomycota. And *Coemansia reversa* (Chang et al., 2015), symbiotic with aquatic arthropods, represents the Zoopagomycota.

By far the most widely studied eukaryotic research organism among the fungi is *Saccharomyces cerevisiae*, baker’s or brewer’s yeast. This ascomycete yeast has for decades been a valuable model organism in research areas including eukaryotic genetics, developmental biology, and biochemistry (Goffeau et al., 1996). Much of what is known about protein import into mitochondria originated in studies of the SAM and TOM protein import complexes of the yeast outer mitochondrial membrane. The *Saccharomyces cerevisiae* SAM complex contains, among other proteins, the Sam37 protein that has a degree of homology with the metaxins and metaxin-like proteins. The Sam37 protein possesses GST_N_Metaxin and GST_C_Metaxin protein domains that identify the metaxins and metaxin-like proteins. But the major domains in the Sam37 protein are the Tom37 and Tom37_C domains. Also, there aren’t two metaxin-like proteins, MTXa and MTXb, as found in other fungi and discussed in this article, only Sam37.

For this reason, Sam37 is a protein that has features shared with the metaxins, but is in a separate category and is not an MTXa or MTXb protein. Sam37 proteins are not confined to *S. cerevisiae*, but are present in other yeasts in the same taxonomic family, Saccharomycetaceae, including *Candida glabrata* and *Pichia stipitis* (Dujon et al., 2004).

Although Sam37 is not a metaxin, Sam37 and the vertebrate metaxins have functional homology, since a role for vertebrate metaxins 1 and 2 in protein import into mitochondria has been experimentally demonstrated. The Tom37 domain found in yeast protein Sam37 is also a major protein domain of the metaxins and metaxin-like proteins, in addition to the GST_N_ and GST_C_Metaxin domains, as shown in this report. The Sam37 protein is part of the SAM complex (<u>S</u>orting and <u>A</u>ssembly <u>M</u>achinery complex), which is made up of Sam37 (previously Tom37), Sam50, Sam35, and Mdm10 proteins. Located in the outer mitochondrial membrane, the SAM complex works with the TOM complex to bring about the uptake and insertion of β-barrel membrane proteins into the outer mitochondrial membrane. Although the details of protein import into vertebrate mitochondria are less well known, the yeast results strengthen the evidence of a role for the metaxin proteins.

The fungal MTXa and MTXb proteins in this study are also compared to fungal FAXC proteins, in addition to vertebrate metaxins. The failed axon connections (*fax*) gene was initially identified in *Drosophila* as a gene that, when mutated, enhanced the mutant phenotype of the *Abl* oncogene (Hill et al., 1995). A human ortholog, the FAXC gene, has been detected at chromosome band 6q16.2. Genes homologous to the FAXC gene exist in many vertebrate and invertebrate species, as well as in fungi. The FAXC protein in humans may have a role in axonal development, which is therefore a different role than the metaxins. Of relevance to this report, the FAXC protein has characteristics of a metaxin-like protein (Adolph, unpublished). The FAXC protein has GST_N_Metaxin and GST_C_Metaxin domains, it has a pattern of predicted α-helices similar to the vertebrate metaxins, and it is phylogenetically related to the metaxins. Because the FAXC protein is structurally related to the metaxins, it has been included in some of the analyses discussed in this paper.

The GST_N_Metaxin and GST_C_Metaxin domains of the metaxins and metaxin-like proteins are distinct from the GST domains found in other glutathione S-transferase superfamily proteins. As examples, among the human glutathione S-transferase proteins, GST A1 has GST_N_Alpha and GST_C_Alpha domains as the defining domains. For GST M1, GST_N_Mu and GST_C_Mu are the major domains. And GST T1 has GST_N_Theta and GST_C_Theta domains. And although GSTs have well-characterized roles in cellular detoxification reactions, most GST protein sequences added to databases represent GST superfamily proteins of unknown function. The vertebrate metaxins are among the GST superfamily proteins with known functions but non-typical GST functions, specifically in protein import into mitochondria.

A previous publication (Adolph, 2021) reported the existence of metaxin-like proteins in fungi representing major fungal taxonomic divisions, most commonly the Mucoromycota and Ascomycota divisions. For these fungi, a single metaxin-like protein was usually detected, with the GST_N_Metaxin, GST_C_Metaxin, and Tom37 domains that identify, along with other criteria, the protein as metaxin-like. Vertebrates and invertebrates possess multiple metaxins, specifically metaxins 1, 2, and 3 in vertebrates and metaxins 1 and 2 in invertebrates. Therefore, the possible existence of multiple metaxin-like proteins in fungi has been investigated further. It was found that two metaxin-like proteins, designated MTXa and MTXb, are present in a variety of fungi. But the MTX proteins are not direct homologs of vertebrate metaxins 1, 2, or 3, since the fungal proteins are about equally homologous to each of the vertebrate metaxins. However, similar to vertebrate metaxins, the fungal metaxin-like proteins MTXa and MTXb represent distinct categories of proteins with little homology between them. The results of this study provide the basis for the further investigation of the roles of the fungal metaxin-like proteins in the process of protein import into mitochondria or other cellular processes.

## 2. METHODS

Identities and similarities of two protein sequences, as in Figure 1, were determined with the NCBI Global Alignment tool (Needleman and Wunsch, 1970; Altschul et al., 1990), available at https://blast.ncbi.nlm.nih.gov/Blast.cgi. The alignment of two protein sequences to compare the highest homology regions used Align Two Sequences BLAST. The EMBOSS Needle tool (https://www.ebi.ac.uk/Tools/psa/emboss_needle/) was also employed to align pairs of sequences. The presence of GST_N_Metaxin, GST_C_Metaxin, and Tom37 domains was investigated with the NCBI CD_Search tool at www.ncbi.nlm.nih.gov/Structure/cdd/wrpsb.cgi (Lu et al., 2020). Figures 2, 3A, and 3B include these conserved domains for the metaxin-like proteins of representative fungi and other organisms. The BLAST Expect (E) values reported for the domains are a measure of the significance of the results of the conserved domain database searches. Low E-values mean more significant domains. Multiple sequence alignments of fungal metaxin-like proteins and other proteins (Figures 4, 5, and 6) were obtained with the COBALT tool of the NCBI (Papadopoulos and Agarwala, 2007), at www.ncbi.nlm.nih.gov/tools/cobalt. Alpha-helical segments were predicted using the PSIPRED server (Jones, 1999; bioinf.cs.ucl.ac.uk/psipred/). Phylogenetic trees, such as those in Figures 7 and 8, were also obtained from the NCBI COBALT multiple sequence alignments. In addition, Clustal Omega was employed (Sievers et al., 2011; https://www.ebi.ac.uk/Tools/msa/clustalo/). Two different servers were used to identify possible transmembrane helices: the TMHMM server (Krogh et al., 2001) at www.cbs.dtu.dk/services/TMHMM/ and the PHOBIUS server (Kall et al., 2007) at http://phobius.sbc.su.se/.

**Figure 1.**
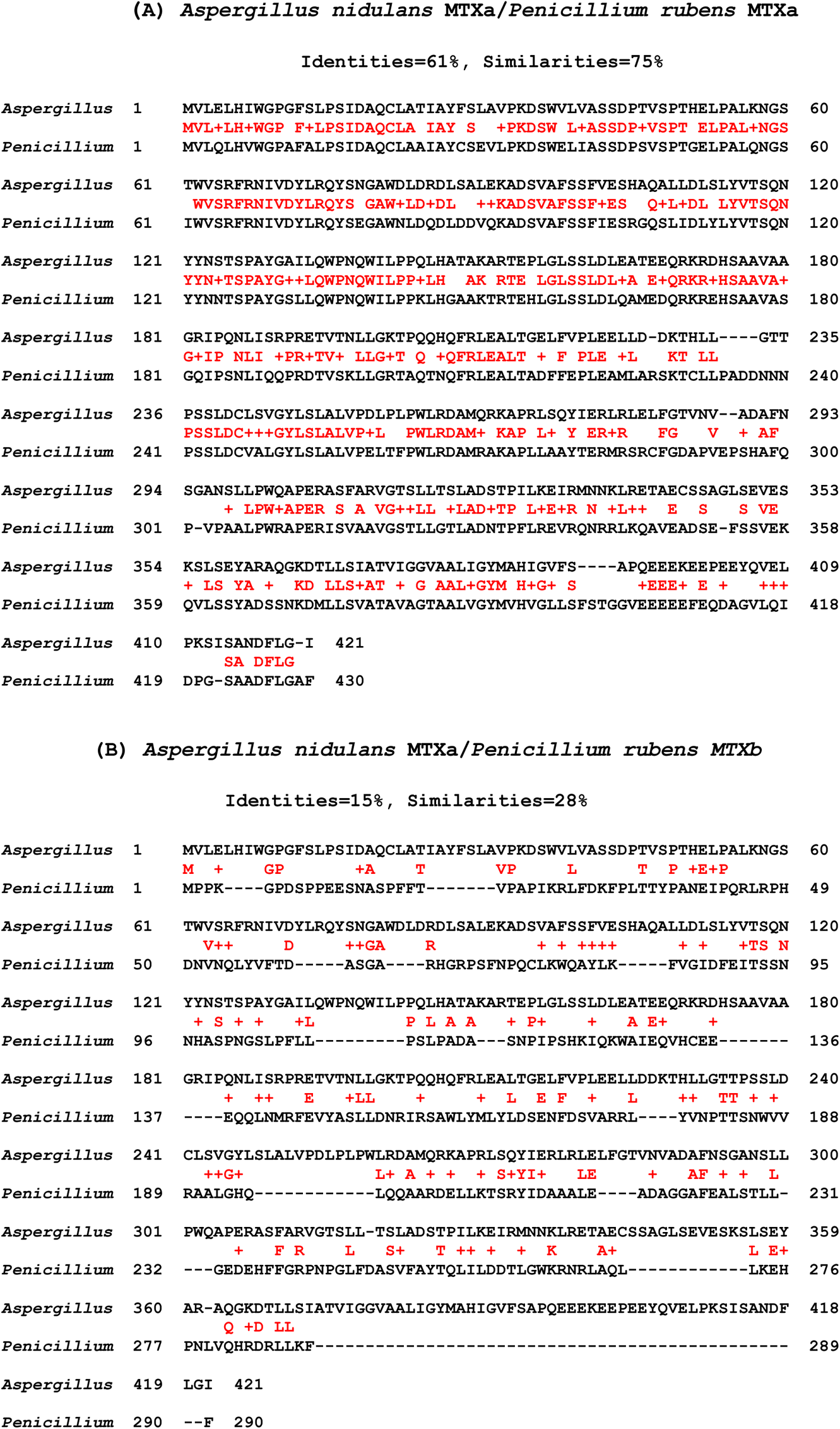
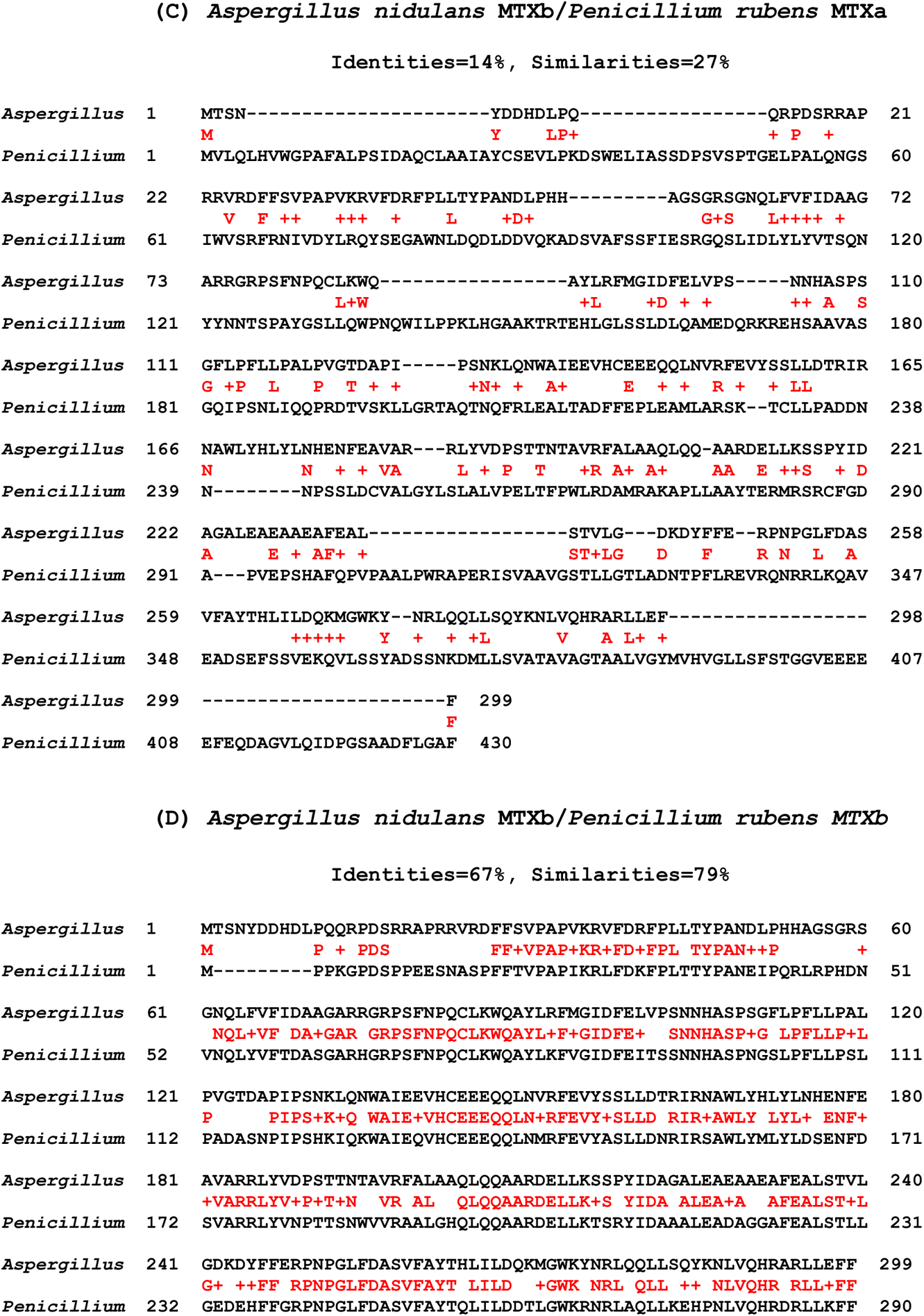
Amino acid sequence identities of MTXa and MTXb proteins of fungi. (A, B) In Figure 1A, the MTXa protein of *Aspergillus nidulans* is aligned with the MTXa protein of *Penicillium rubens*. Identical amino acids are shown in red between the aligned sequences, with similar amino acids indicated by “+” signs. The two fungi are in the Ascomycota division. In Figure 1B, *Aspergillus nidulans* MTXa is aligned with *Penicillium rubens* MTXb. The percentages of amino acid identities are strikingly different in (A) with 61% identities and in (B) with only 15%. These and other results show convincingly that the MTXa and MTXb fungal proteins are separate and distinct categories. *Aspergillus nidulans* MTXa and MTXb have 421 and 299 amino acids, respectively, while *Penicillium rubens* MTXa and MTXb have 430 and 290. The sequences were aligned using NCBI BLAST Global Align. (C, D) Comparable results to those in Figure 1A and 1B are found in aligning *Aspergillus nidulans* MTXb with *Penicillium rubens* MTXa and MTXb. With MTXa of *P. rubens*, the identities are only 14% (Figure 1C), while with *P. rubens* MTXb, 67% identities are found (Figure 1D). The high degree of homology shown in (A) and (D) means that the MTXa proteins have a close evolutionary relationship, as do the MTXb proteins. But the low percentages in (B) and (C) mean there is little relationship between the MTXa and b proteins, as detected by amino acid alignments. A low degree of homology is also seen in alignments of MTXa proteins with MTXb proteins of the same fungi. Alignment of MTXa and MTXb of *A. nidulans* reveals only 15% amino acid identities. *P. rubens* MTXa and MTXb show only 17% identities.

**Figure 2.**
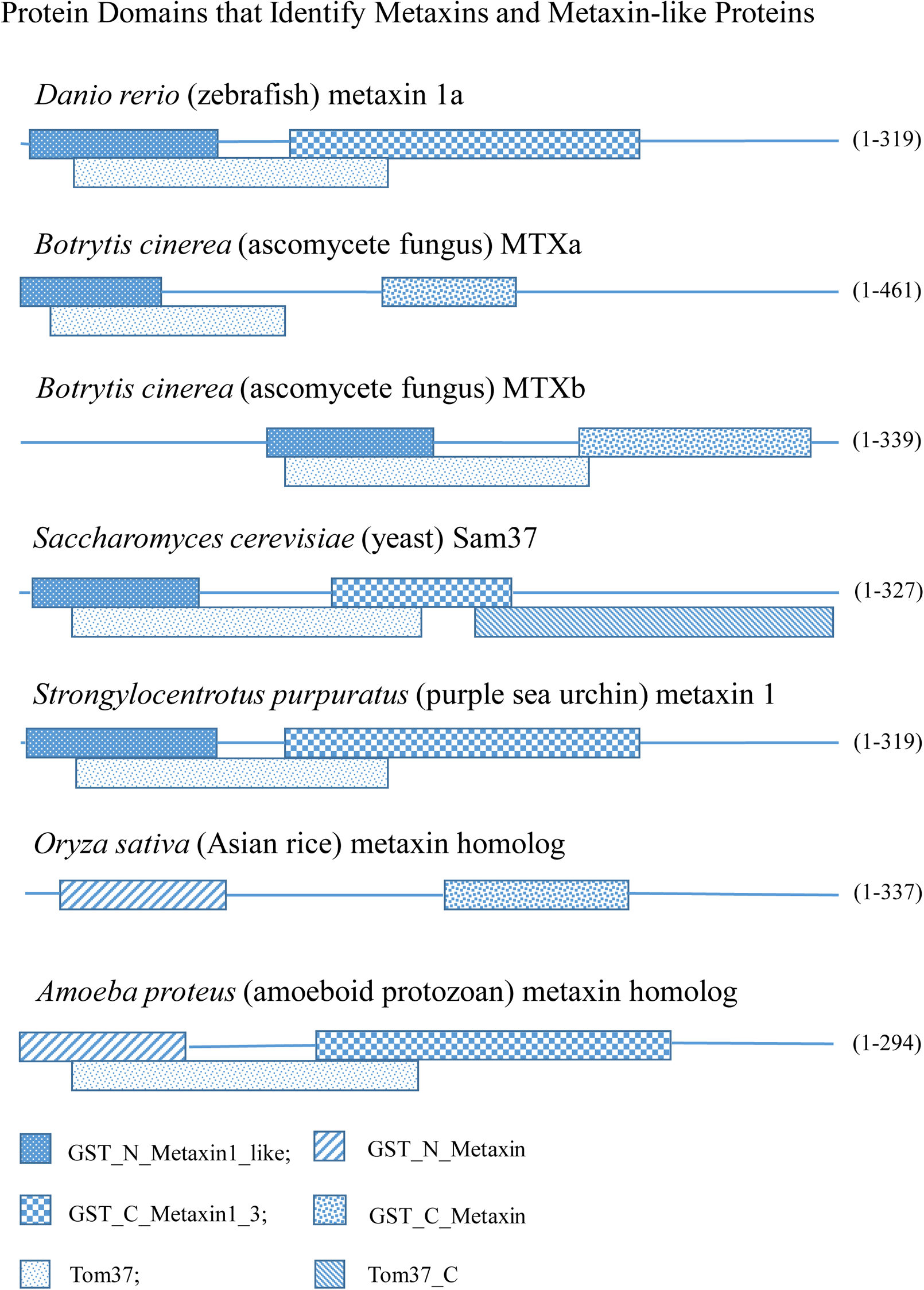
Protein domains of metaxins, metaxin-like proteins, and Sam37. The figure shows the domain structures of the metaxin proteins or metaxin-like proteins of a representative vertebrate, fungus, yeast, invertebrate, plant, and protist. The GST_N_Metaxin and GST_C_Metaxin domains that define proteins as being metaxins or metaxin-like are highly conserved in these examples. Zebrafish (*Danio rerio*) metaxin 1a has metaxin domains typical of vertebrates, with BLAST Expect (E) values characteristic of major domains. The wine grape fungus *Botrytis cinerea* has multiple metaxin-like proteins, MTXa and MTXb, like other fungi as demonstrated in this paper. For the Sam37 protein of the yeast *Saccharomyces cerevisiae*, the Tom37 and Tom37_C domains are the major, defining protein domains. But GST_N_Metaxin and GST_C_Metaxin domains are also prominent. Tom37 is a domain associated with the Sam37 protein, and is also present in the vertebrate (zebrafish) and invertebrate (sea urchin) metaxins in Figure 2, and in the fungal, plant, and protist metaxin-like proteins. The sea urchin *Strongylocentrotus purpuratus* has a metaxin 1 protein with major GST_N_Metaxin and GST_C_Metaxin domains, similar to vertebrates. The sea urchin also has a metaxin 2 protein (not shown). The plant *Oryza sativa* (cultivated rice) has a single metaxin homolog. A single metaxin-like protein is also found for *Amoeba proteus*, a protist or amoeboid protozoan. In addition, bacteria (not included in the figure) have a single metaxin homolog. All these examples demonstrate the great diversity of organisms that have metaxins or metaxin-like proteins.

**Figure 3.**
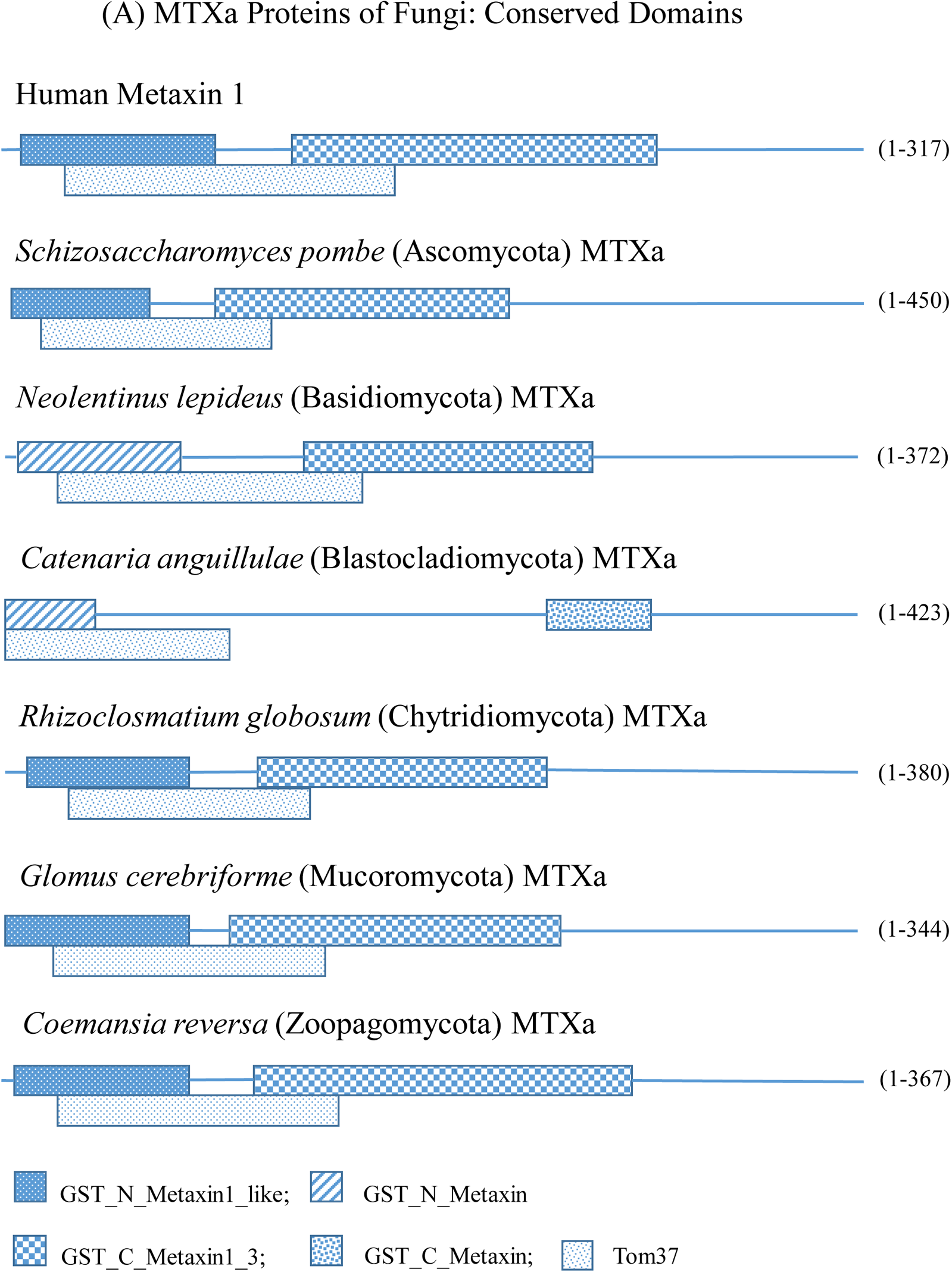
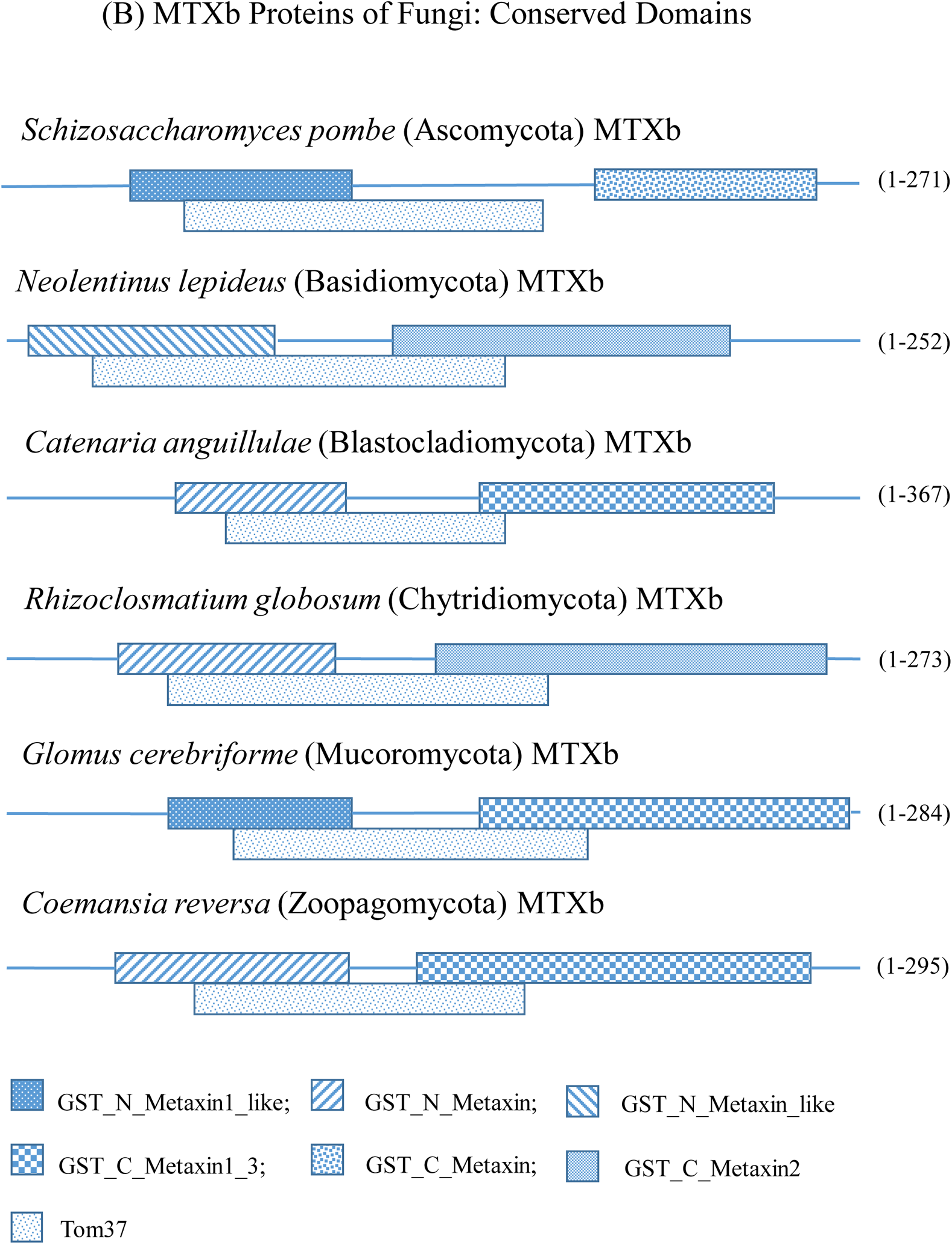
Conserved domains of MTXa and MTXb proteins of fungi. (A) The MTXa domain structures are shown for a selection of fungi of major taxonomic divisions: Ascomycota (*Schizosaccharomyces pombe*), Basidiomycota (*Neolentinus lepideus*), Blastocladiomycota (*Catenaria anguillulae*), Chytridiomycota (*Rhizoclosmatium globosum*), Mucoromycota (*Glomus cerebriforme*), and Zoopagomycota (*Coemansia reversa*). The human metaxin 1 domains are also included for comparison. GST_N_Metaxin and GST_C_Metaxin domains are present in all the examples shown, as is the Tom37 domain. The domains are generally weaker (i.e., have higher E values) in the fungi than in human metaxin 1. The C-terminal 30–33% of the protein chains are devoid of major domains, and the GST_N_Metaxin domains begin close to the N-terminus. (B) The characteristic domains of the MTXb proteins of the same fungi as in (A) are also the GST_N_Metaxin, GST_C_Metaxin, and Tom37 domains. The MTXb GST_N_Metaxin domains generally begin further from the N-terminus, and the GST_C_Metaxin domains end closer to the C-terminus. For example, the GST_N_Metaxin domain of the MTXa protein of *Schizosaccharomyces pombe* begins at the N-terminus (Figure 3A), while the GST_N_Metaxin domain of the MTXb protein starts 40 bases (15%) from the N-terminus. The MTXa proteins in Figure 3A average 389 amino acids, while the MTXb proteins in Figure 3B average 290 amino acids. The MTXb protein chains are therefore 25% shorter than the MTXa chains.

**Figure 4.**
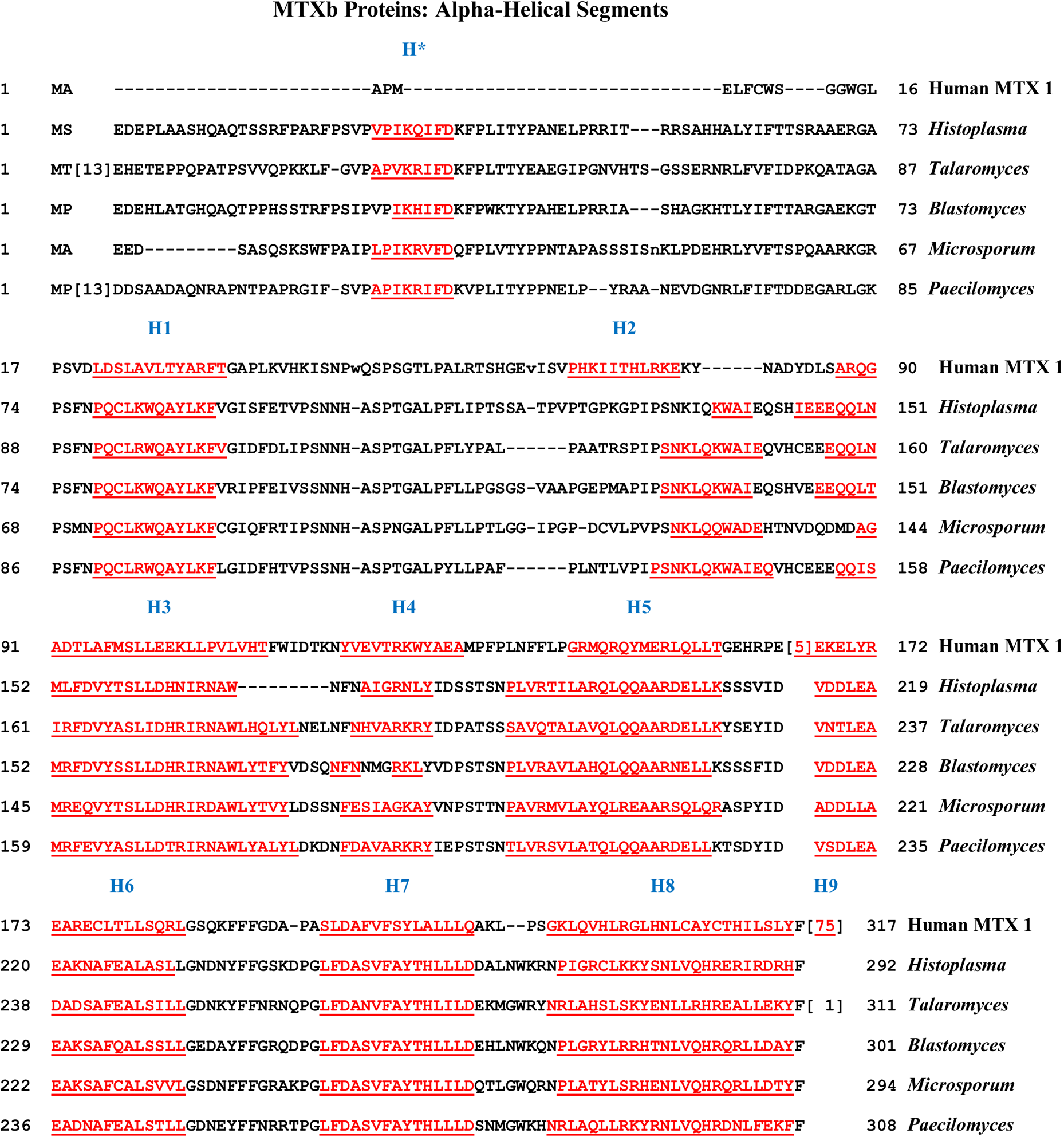
Alpha-helical segments of representative MTXb metaxin-like proteins of fungi. The multiple sequence alignment shows the predicted α-helical segments in red and underlined. The MTXb proteins of five fungi of the Ascomycota division are included: *Histoplasma capsulatum*, *Talaromyces marneffei*, *Blastomyces dermatitidis*, *Microsporum canis*, and *Paecilomyces variotii*. All are potential human pathogens. Also included in the figure is human metaxin 1. The five MTXb proteins all possess helices H1 through H8, as does human metaxin 1. The MTXb proteins have an extra N-terminal helix at position H*, which is absent in human metaxin 1. But the fungal proteins lack the C-terminal helix H9 found in human metaxin 1 (within sequence “<u>75</u>”). Human metaxin 2 also possesses an extra N-terminal helix at H* and lacks C-terminal helix H9. However, amino acid sequence alignments demonstrate that fungal MTXb proteins are not direct homologs of metaxin 2. But, as discussed in section 3.2, MTXb proteins may, to a small degree, be more homologous to vertebrate metaxin 2 proteins, and MTXa proteins to vertebrate metaxin 1 proteins. MTXa proteins of the same fungi in the figure also reveal a conserved pattern of α-helical segments (not shown). In this case, H1 through H9 are present, with the helix at H* absent, similar to human metaxin 1. Amino acid alignments, however, show that MTXa proteins are not simply fungal metaxin 1 homologs (section 3.2). But the resemblance of the patterns of α-helices of the MTXa and MTXb proteins to those of human metaxins 1 and 2, respectively, may indicate, as mentioned above, a degree of homology between MTXa and metaxin 1 proteins, as well as MTXb and metaxin 2 proteins.

**Figure 5.**
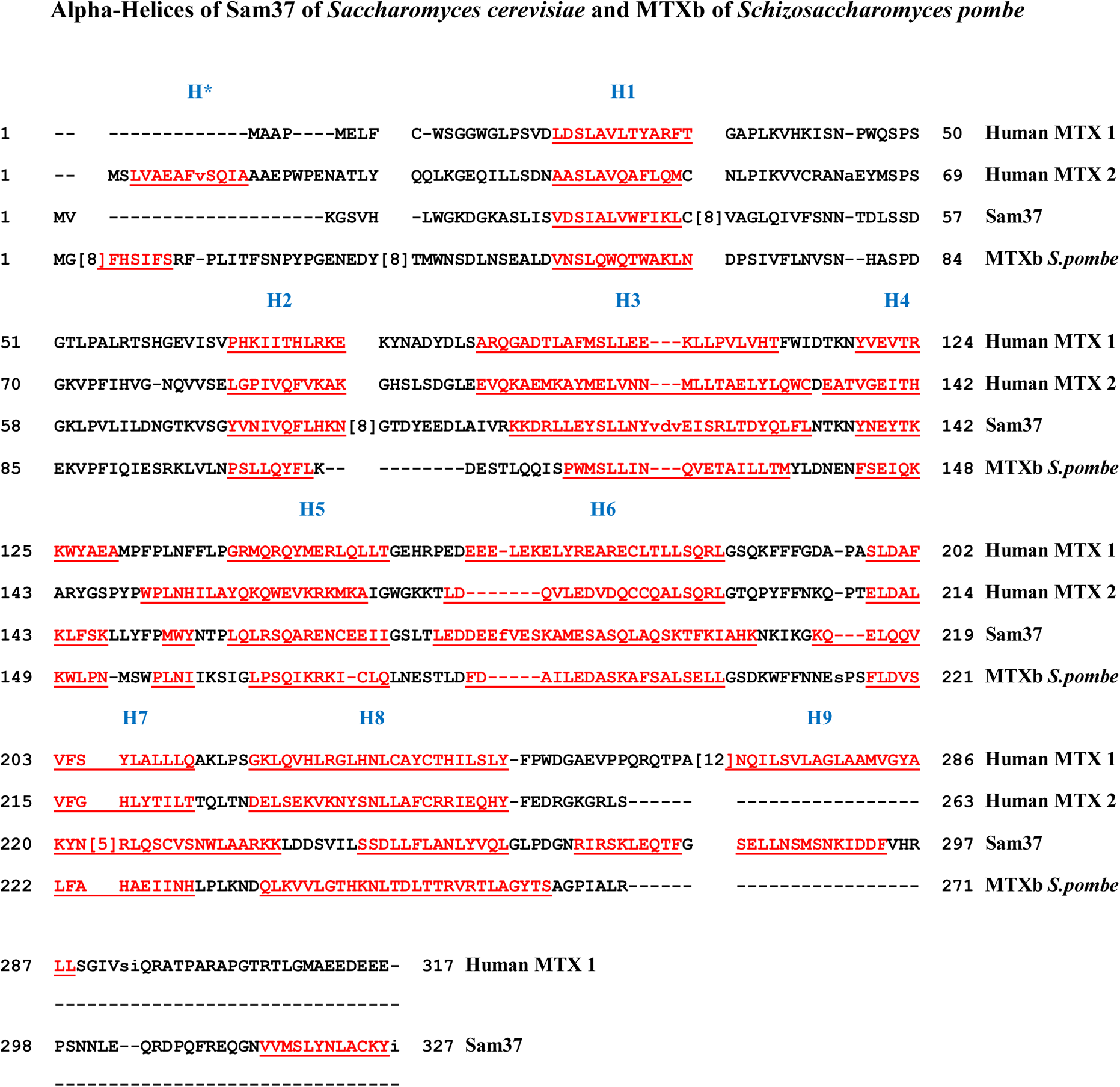
Protein secondary structure of Sam37 and fission yeast MTXb: multiple sequence alignments. In the figure, the α-helical structure of *Saccharomyces cerevisiae* Sam37 is compared to that of *Schizosaccharomyces pombe* MTXb, and also human metaxin 1 and metaxin 2. Sam37 is a protein of the outer mitochondrial membrane of baker’s or brewer’s yeast (*Saccharomyces cerevisiae*), and has a well-characterized role as a component of the SAM complex in the import of proteins into yeast mitochondria. The major domains of *S. cerevisiae* Sam37 are the Tom37 and Tom37_C domains (Figure 2). Sam37 is also a metaxin-related protein because it possesses the GST_N_Metaxin and GST_C_Metaxin domains that characterize the metaxins and metaxin-like proteins. *Schizosaccharomyces pombe* (“fission yeast”) is an important model organism used in cell cycle regulation studies. Like *S. cerevisiae*, it is in the Ascomycota division, but has MTXa and MTXb proteins like fungi of other divisions and unlike *S. cerevisiae* yeast. As Figure 5 demonstrates, Sam37 and fission yeast MTXb are α-helical proteins with similar patterns of helical segments. Helices H1 through H8 are present in all the aligned protein sequences. The high level of conservation of H1 – H8 is a prominent feature of the secondary structures of metaxins and metaxin-like proteins. Sam37 has a pattern of α-helices like that of human metaxin 1. It lacks helix H* found in human metaxin 2, but has an extra C-terminal helix that coincides with H9. *S. pombe* MTXb has a helix that largely coincides with H* near the N-terminus but lacks H9, while MTXa (not shown) is similar to human metaxin 1 with an extra C-terminal helix.

**Figure 6.**
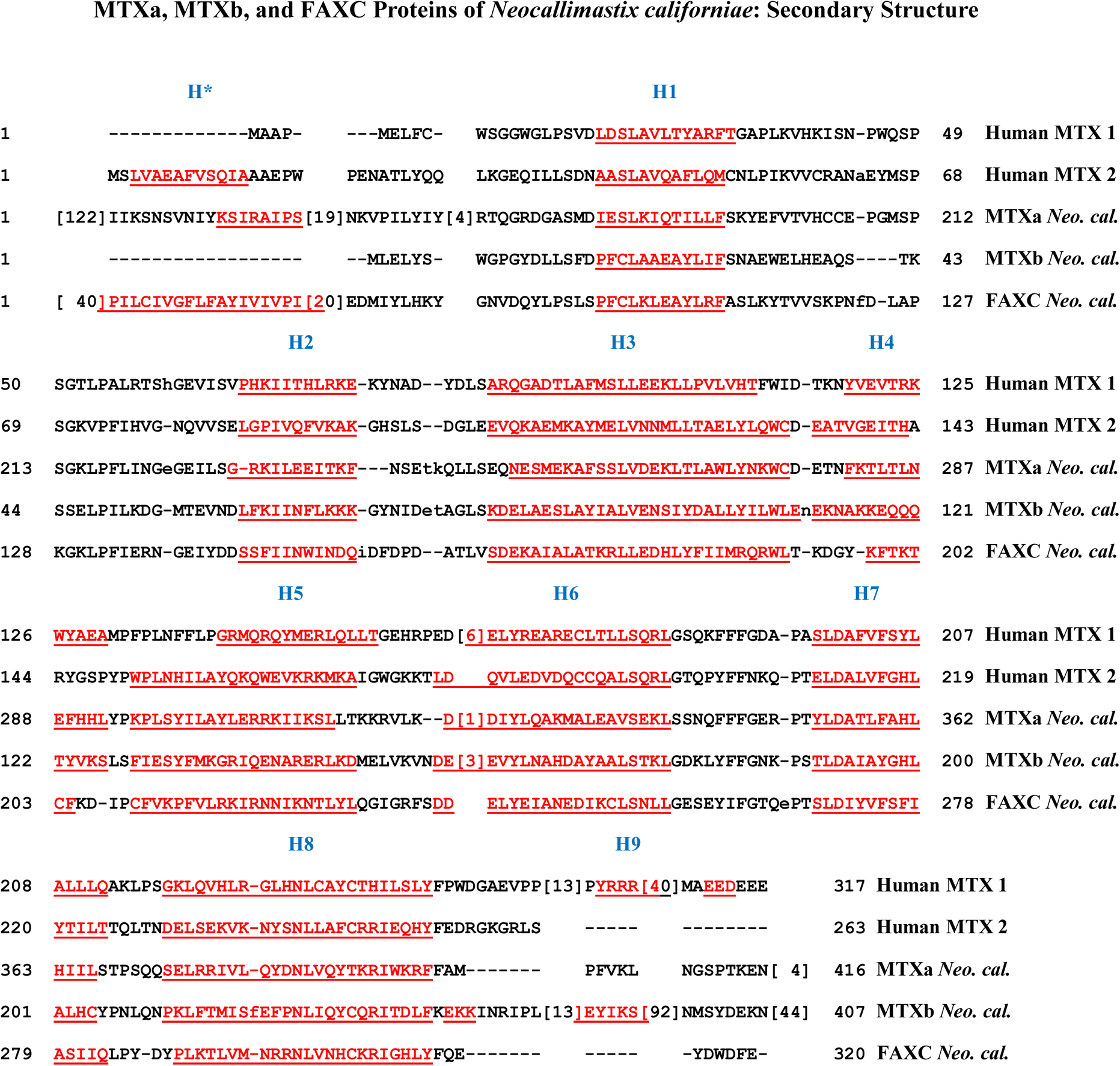
Secondary structure of MTXa, MTXb, and FAXC metaxin-like proteins. The figure shows the highly conserved pattern of α-helical segments in all three distinct metaxin-like proteins from *Neocallimastix californiae* (abbreviated *Neo. cal.*). The fungus is in the Neocallimastigomycota division, and is a microbiome fungus of the digestive tract of large herbivores. In addition to the Neocallimastigomycota division, fungi with MTX and FAXC proteins were found to exist in other divisions, including the Chytridiomycota and Basidiomycota divisions. The pattern of α-helical segments for all three metaxin-like proteins clearly resembles that in Figure 4 and Figure 5. Eight helices, H1 through H8, are present in each of the fungal proteins, and also in human metaxin 1 and metaxin 2. Extra N-terminal helices (near position H*) are in human metaxin 2 and in MTXa and FAXC. C-terminal helix H9 is in human metaxin 1, and an extra C-terminal helix is also in MTXb.

**Figure 7.**
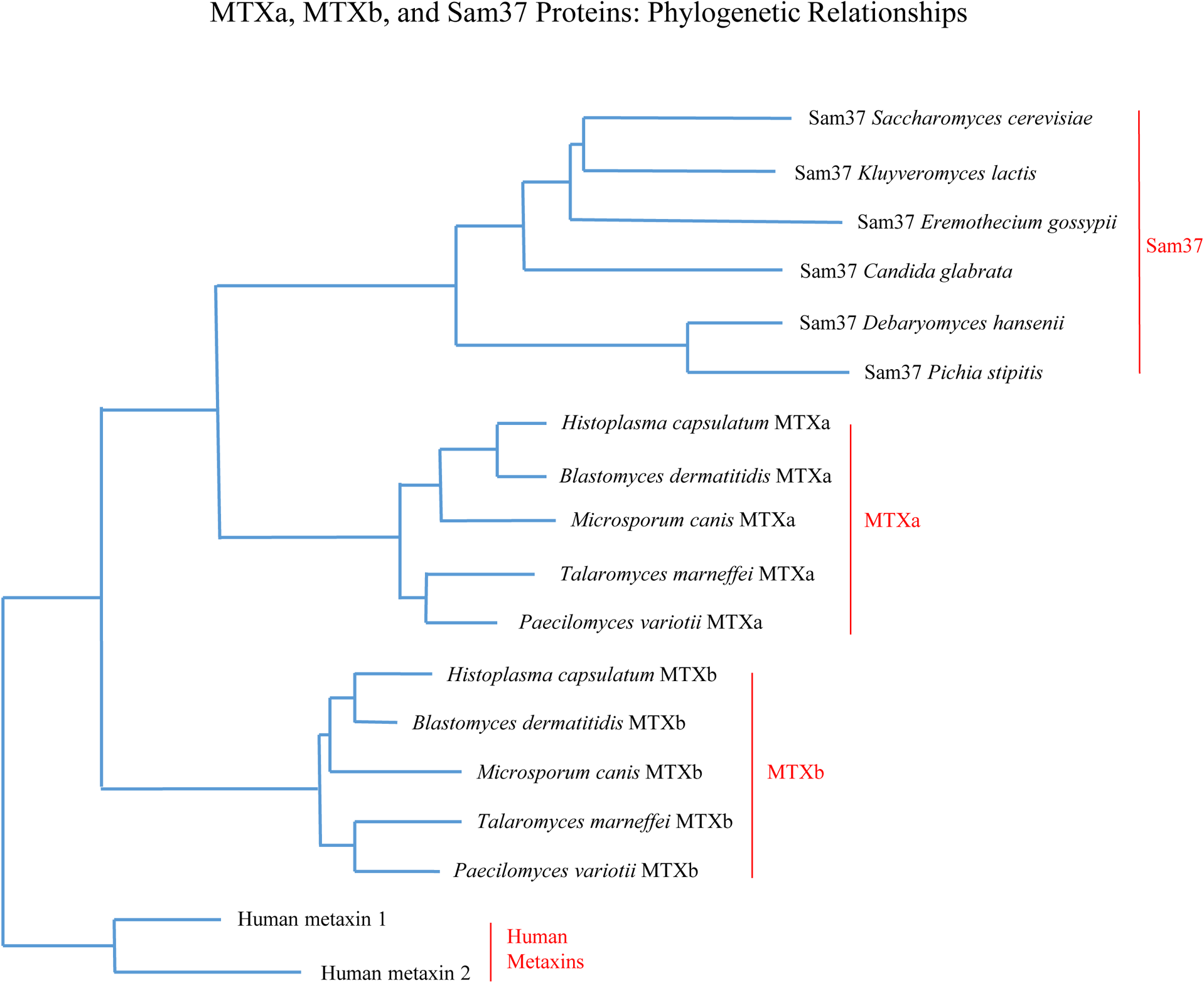
Phylogenetic relationships of MTXa, MTXb, and Sam37 proteins. The MTXa and MTXb proteins in the figure are those of fungal species in the Ascomycota division. These include the potential human pathogens *Histoplasma capsulatum, Blastomyces dermatitidis, Microsporum canis, Talaromyces marneffei,* and *Paecilomyces variotii*. Although the MTXa and MTXb proteins are alike in possessing GST_N_Metaxin and GST_C_Metaxin domains, the phylogenetic analysis shows that MTXa proteins form a separate group, as do MTXb proteins. These results clearly demonstrate that MTXa proteins and MTXb proteins are distinct categories of metaxin-like proteins. *Saccharomyces cerevisiae* Sam37 protein has GST_N_ and GST_C_Metaxin domains, and also major Tom37 and Tom37_C domains. But *S. cerevisiae* Sam37 is in a separate category from the fungal MTXa or MTXb proteins, as the phylogenetic analysis indicates. The same is true for five other Sam37 proteins of Saccharomycetaceae family yeasts or fungi in Figure 7: *Kluyveromyces lactis, Eremothecium gossypii, Candida glabrata*, *Debaryomyces hansenii*, and *Pichia stipitis*. The Sam37 proteins form a distinct and separate group of metaxin-related proteins. In addition, the figure shows that human metaxins 1 and 2 are related by evolution to fungal metaxin-like proteins MTXa and MTXb, as well as SAM37 proteins, and share a common ancestral protein sequence. These phylogenetic relationships reflect the features shared by human metaxins, fungal metaxin-like proteins, and SAM37s, including GST_N_Metaxin and GST_C_Metaxin domains and similar conserved patterns of α-helical segments.

**Figure 8.**
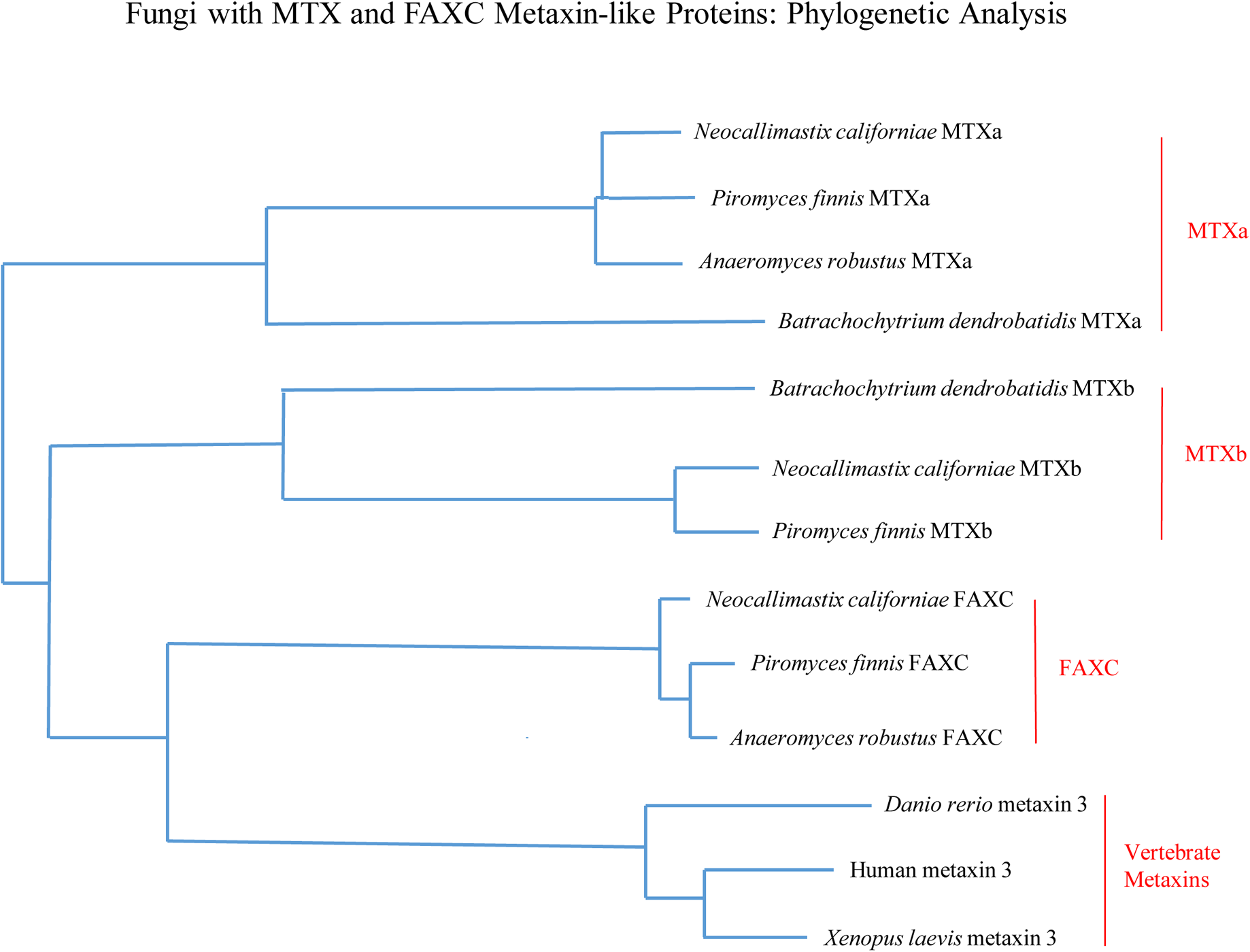
Evolutionary relationships of MTXa, MTXb, and FAXC proteins. In fungi with multiple metaxin-like proteins, having two proteins, MTXa and MTXb, is the most common. But numerous fungi also possess metaxin-like FAXC proteins. The three metaxin-like proteins, MTXa, MTXb, and FAXC, are found in fungi of divisions that include the Neocallimastigomycota, Chytridiomycota, and Basidiomycota divisions. In Figure 8, *Neocallimastix californiae* and *Piromyces finnis* are in the Neocallimastigomycota division. Two additional fungi are included in the figure, *Anaeromyces robustus* (Neocallimastigomycota) with MTXa and FAXC proteins only, and *Batrachochytrium dendrobatidis* (Chytridiomycota) with MTXa and b proteins only. The vertebrate metaxin 3 proteins in the figure – human, *Danio rerio* (zebrafish), and *Xenopus laevis* (frog) – reveal that vertebrate metaxins are related by evolution to the fungal metaxin-like proteins. Vertebrate metaxins are also related to the metaxins and metaxin-like proteins of invertebrates, plants, bacteria, and protists, as discussed in the Introduction.

## 3. RESULTS AND DISCUSSION

### 3.1. Identification of Multiple Metaxin-like Proteins in Fungi

The presence of multiple metaxin-like proteins in fungi was investigated by searches of protein sequence databases of the NCBI and JGI (DOE Joint Genome Institute). Metaxin-like proteins were identified by criteria including homology with vertebrate metaxins and the possession of characteristic GST_N_Metaxin and GST_C_Metaxin protein domains. The Tom37 conserved protein domain was frequently present, but not other major, unrelated protein domains.

Fungi with two distinct metaxin-like proteins, MTXa and MTXb, were identified in major taxonomic divisions (phyla) including Ascomycota, Basidiomycota, Blastocladiomycota, Chytridiomycota, Mucoromycota, Neocallimastigomycota, and Zoopagomycota. The divisions with the greatest number of metaxin-like proteins in this study were Ascomycota and Mucoromycota. The ascomycete fungi include the model organisms *Saccharomyces cerevisiae*, baker’s or brewer’s yeast, and *Schizosaccharomyces pombe*, fission yeast. The MTXa proteins of different fungi form one distinct category, while the MTXb proteins form another. MTXa and MTXb are not directly homologous to vertebrate metaxin 1 and metaxin 2, respectively. Instead, MTXa is about equally homologous to each of the vertebrate metaxins, and the same is true for MTXb.

FAXC metaxin-like proteins, in addition to MTX proteins, were found to be present in fungi of different taxonomic divisions. FAXC is, like the fungal MTXa and MTXb proteins, about equally homologous to vertebrate metaxins 1, 2, and 3. MTXa and b proteins and FAXC proteins constitute three distinct categories of metaxin-like proteins, as shown by amino acid sequence alignments, discussed in section 3.2, and phylogenetic analysis, discussed in section 3.6.

### 3.2. MTXa and MTXb Proteins of Fungi: Amino Acid Sequence Alignments

Amino acid sequence alignments show that MTXa and MTXb proteins are distinct categories of fungal metaxin-like proteins. Examples are included in Figure 1. Alignment of MTXa of *Aspergillus nidulans* with MTXa of *Penicillium rubens* reveals a high degree of homology in the amino acid sequences, with 61% identities and 75% similarities (Figure 1A). *Aspergillus nidulans* (Galagan et al., 2005) is a widely used model organism in eukaryotic cell biology research, while *Penicillium rubens* (van den Berg et al., 2008) is an important source of the antibiotic penicillin. They are both in the Ascomycota division. A major difference is seen in aligning *Aspergillus nidulans* MTXa and *Penicillium rubens* MTXb, which shows only 15% amino acid identities (Figure 1B). An analogous result is encountered in aligning *Aspergillus nidulans* MTXb and *Penicillium rubens* MTXa, with only 14% identities. But 67% identities are found in aligning the MTXb protein of *A. nidulans* and the MTXb protein of *P. rubens* (Figure 1C, D). In addition, MTXa and MTXb proteins of *A. nidulans* show only 15% identities when aligned with each other, and MTXa and MTXb proteins of *P. rubens* only 17%.

Alignments of other MTXa and MTXb proteins show similar results. For the MTXa proteins of *Fusarium verticillioides* (Ascomycota), a destructive fungus of maize, and *Trichoderma virens* (also Ascomycota), a beneficial symbiont of plants, 62% identities are found. But the identities decrease to 16% for MTXa of *F. verticillioides* and MTXb of *T. virens*. The MTXa proteins of *Absidia repens* and *Hesseltinella vesiculosa* (both Mucoromycota) show 65% identities. But when the *H. vesiculosa* protein is MTXb, the identities decrease to 28%. A third example is the MTXa and MTXb proteins of the Basidiomycota fungi *Heliocybe sulcata* and *Neolentinus lepideus*. 74% identities result when the MTXa proteins are aligned, but only 17% when the *N. lepideus* protein is MTXb.

All these and other amino acid alignment results clearly demonstrate that MTXa and MTXb proteins are distinct categories of fungal metaxin-like proteins. It is therefore not only vertebrate and invertebrate metaxins that exist as more than a single protein species.

Are the fungal MTXa and MTXb proteins direct homologs of vertebrate metaxin 1 and metaxin 2? Amino acid sequence alignments show that MTXa and MTXb are about equally homologous to vertebrate metaxins 1 and 2. However, some alignments suggest that MTXa has a slightly higher degree of homology to vertebrate metaxin 1, and MTXb to vertebrate metaxin 2. An example that shows equal homology is that of *Absidia repens* (Mucoromycota), a fungus of decaying plant matter. In this case, *A. repens* MTXa has 24% amino acid identities with human metaxin 1 and also 24% with human metaxin 2. MTXb shows 29% identities with both human metaxins 1 and 2. A different conclusion than equal homology might be drawn from the example of *Aspergillus fumigatus* (Ascomycota), a cause of fungal infections (aspergillosis) in humans. Here, MTXa and human metaxin 1 have 26% identities, but MTXa and human metaxin 2 have “no significant similarity”. However, the inherent imprecision in the percent identities of the amino acid sequence alignments is too large to allow the conclusion that, for *A*. *fumigatus*, MTXa is a fungal metaxin 1 and, similarly, MTXb is a fungal metaxin 2. This imprecision exists using both BLAST Align Two Sequences and also Global Align, which give comparable but not exactly identical results.

For fungi with both MTX and FAXC proteins, the two types of proteins have a similar degree of homology to each of the vertebrate metaxins, again taking into account the inaccuracies in the percent identities. However, the MTX proteins and FAXC proteins are distinct categories of proteins, as shown by amino acid sequence alignments. For example, alignment of *Neocallimastix californiae* MTXa with *N*. *californiae* FAXC reveals just 18% identities, and MTXb with FAXC, 14%. These distinct MTX and FAXC proteins are conserved in other closely related fungi. For example, *Neocallimastix californiae* and *Piromyces finnis* are in the same taxonomic division, class, order, and family. They also have highly conserved MTXa, MTXb, and FAXC proteins. Alignment of *N. californiae* MTXa and *P. finnis* MTXa reveals 74% amino acid identities, *N. californiae* MTXb with *P. finnis* MTXb reveals 77% identities, and *N. californiae* FAXC with *P. finnis* FAXC, 84%. But for *N. californiae* MTXa and *P. finnis* FAXC, the percentage drops to 19%, and low percentages are also found with other, similar alignments.

### 3.3. Presence of Metaxin Protein Domains in the Multiple Metaxin-like Proteins of Fungi

A defining characteristic of metaxin-like proteins is the presence of conserved protein domains. These are the GST_N_Metaxin and GST_C_Metaxin domains initially identified in vertebrate metaxins. Also commonly found is the Tom37 domain, the major protein domain of the Sam37 protein of yeast (*Saccharomyces cerevisiae*). Figure 2 includes the GST_N_Metaxin, GST_C_Metaxin, and Tom 37 domains of metaxin proteins or metaxin-like proteins in a representative vertebrate, fungus, yeast, invertebrate, plant, and protist. The domains are highly conserved, although their strength (BLAST E value), size, and location along the amino acid chains show some differences.

The metaxins were originally described in vertebrates, in particular the mouse, where two metaxin proteins, metaxins 1 and 2, were identified. The proteins were found to be widespread in vertebrates. In Figure 2, zebrafish (*Danio rerio*) metaxin 1a is an example (zebrafish genome sequence: Howe et al., 2013). The protein domain structure of zebrafish metaxin 1a is similar to the structure of mouse metaxin 1, human metaxin 1, and other vertebrate metaxins, with GST_N_Metaxin (E-value: 1.27e-28), GST_C_Metaxin (E-value: 1.12e-76), and Tom37 domains. Fungal metaxin-like proteins are represented in Figure 2 by *Botrytis cinerea* MTXa and MTXb, two distinct metaxin-like proteins. *Botrytis cinerea* is of economic importance as a phytopathogenic fungus that infects wine grapes producing “gray rot” and “noble rot”. Its genome has been sequenced (Anselem et al., 2011). The domain structure of yeast Sam37 shown in Figure 2 is dominated by Tom37 (E-value: 1.29e-39) and Tom37_C (4.85e-72) domains. Sam37 also has GST_N_Metaxin and GST_C_Metaxin domains, a feature shared with the metaxins and metaxin-like proteins in the figure.

Invertebrates such as the sea urchin *Strongylocentrotus purpuratus* in Figure 2 possess metaxin 1 and metaxin 2 proteins with domain structures similar to the vertebrate metaxins. Metaxin 1 of the sea urchin, which has a sequenced genome (Sodergren et al., 2006), shares a high percentage of amino acid identities (44%) with human metaxin 1 and has strong, vertebrate-like GST_N_Metaxin (E-value: 4.48e-30) and GST_C_Metaxin (E-value: 3.84e-78) domains. In contrast to vertebrates and invertebrates with distinct metaxin 1 and 2 proteins, plants and protists have only a single metaxin-like protein. As examples, Figure 2 includes the domain structures of the metaxin-like proteins of Asian rice (*Oryza sativa*; genome sequence: Goff et al., 2002) and an amoeboid protist (*Amoeba proteus*).

In Figure 3, conserved protein domains are shown for the metaxin-like proteins MTXa and MTXb of representative fungal species of major taxonomic divisions. The fungi include *Schizosaccharomyces pombe* (Ascomycota), a model eukaryote in biology research, *Neolentinus lepideus*, a basidiomycete mushroom, *Catenaria anguillulae*, a parasitic fungus of the Blastocladiomycota, *Rhizoclosmatium globosum*, a flagellated Chytridiomycota fungus, *Glomus cerebriforme*, a Mucoromycota fungus that forms symbiotic relationships (mycorrhizas) with plant roots, and *Coemansia reversa*, a member of the Zoopagomycota.

The MTXa proteins in Figure 3A and the MTXb proteins in Figure 3B both have the GST_N_Metaxin and GST_C_Metaxin domains that define these proteins as metaxin-like. MTXb proteins are typically shorter than the MTXa proteins, with amino acid chains that are about 75% of the MTXa length. The MTXa domains generally have lower (i.e., more significant) E-values. For the example in Figure 3 of *Glomus cerebriforme*, the E-values for the GST_N_ and GST_C_ Metaxin domains of MTXa are 4.16e-22 and 1.87e-24, respectively, and 6.61e-13 and 3.67e-16 for MTXb. Fungal metaxin-like protein domains generally have higher, less-significant E-values compared to the GST_N_ and GST_C_Metaxin domains of vertebrates, which can be as low as the e-70 range.

MTXb proteins of Ascomycota fungi, but not fungi of other taxonomic divisions, typically have a weak Sam35 domain, in addition to major GST_N_Metaxin and GST_C_Metaxin domains. Sam35 is a protein of the SAM complex in yeast that works with the TOM complex in the insertion of β-barrel membrane proteins into the outer mitochondrial membrane. The MTXb protein of *Aspergillus fumigatus*, a cause of invasive fungal infections (aspergillosis), is an example. The GST_N_Metaxin domain (E-value: 5.36e-20) and GST_C_Metaxin domain (E-value: 9.96e-11) identify the protein as metaxin-like. The Sam35 domain is less significant (E-value: 1.94e-06) and shorter, with only about 20 amino acids that are homologous to the Sam35 consensus sequence. In addition, BLAST Global Align showed only 20% amino acid identities in aligning the sequences of *A. fumigatus* MTXb and yeast Sam35. Further, BLAST Align Two Sequences had a query coverage of just 14% in aligning the amino acid sequences. Protein BLAST searches identify the *A. fumigatus* MTXb protein as a putative Sam35 protein. But the results presented here, and the results for other fungi, indicate that the Ascomycota MTXb proteins are not the same as yeast Sam35 proteins. The MTXb proteins can be described as metaxin-like proteins with GST_N_Metaxin and GST_C_Metaxin domains and with weak (hig h E-value) Sam35 domains.

### 3.4. Secondary Structures of the Multiple Metaxin-like Proteins of Fungi: Alignments of α-Helical Segments

Figure 4 includes a multiple-sequence alignment of the MTXb proteins of five Ascomycota fungi. The predicted α-helical segments in each of the amino acid sequences are shown in red and underlined. As the figure demonstrates, the secondary structures of fungal metaxin-like proteins are dominated by α-helices. Extensive β-strand segments are not seen. The five fungi in the figure are *Histoplasma capsulatum*, a common cause of fungal respiratory infections (histoplasmosis), *Talaromyces marneffei*, a fungal pathogen that can produce fatal infections (talaromycosis), *Blastomyces dermatitidis*, the causative agent of the serious fungal infection blastomycosis, *Microsporum canis*, a common source of fungal head infections in humans, and *Paecilomyces variotii*, an opportunistic human pathogen in immunocompromised individuals.

The patterns of α-helical segments are strikingly similar for human metaxin 1 and all five MTXb proteins. Helices H1 through H8 are present in all proteins, and the spacings between the segments are almost identical. Human metaxin 1 has an extra helix, H9, near the C-terminus, while human metaxin 2 (not shown) has an extra N-terminal helix H* and lacks helix H9. The fungal MTXb proteins also have an extra helix near the N-terminus that coincides with H* and, in addition, lack helix H9. However, amino acid sequence alignments (see section 3.2) show that MTXa proteins and MTXb proteins have about equal homology to both human metaxin 1 and human metaxin 2. Therefore, MTXb proteins are not simple homologs of vertebrate metaxin 2. But whether MTXb proteins are somewhat more homologous to vertebrate metaxin 2 proteins, and MTXa proteins to vertebrate metaxin 1 proteins, cannot be ruled out, given the inherent inaccuracies in the percent amino acid identities in the sequence alignments.

The same general pattern of α-helical segments as shown in Figure 4 for MTXb proteins of Ascomycota fungi is also found for the MTXa proteins of the same fungi. There are some differences, such as the absence of helix H* for the MTXa proteins but the presence of a C-terminal helix that coincides with H9. The results are not limited to these Ascomycota fungi. MTXa and MTXb proteins of other major taxonomic divisions of fungi have similar conserved patterns of α-helical segments: MTXa with helix H1 through helix H9, and MTXb with helix H* through helix H8. These results are found for MTXa and MTXb proteins of fungi in taxonomic divisions that include Basidiomycota, Blastocladiomycota, Chytridiomycota, Mucoromycota, Neocallimastigomycota, and Zoopagomycota.

Invertebrates such as the sea urchin, horseshoe crab, and starfish, as well as insects, have metaxin 1 proteins and metaxin 2 proteins like vertebrates. The predicted secondary structures of invertebrate metaxins 1 and 2 have the same patterns of α-helical segments as the human metaxins (Adolph, 2020a). Eight helical segments are found, with an extra C-terminal helix for the invertebrate metaxin 1 proteins and an extra N-terminal helix for the invertebrate metaxin 2 proteins. Plants and bacteria have a single metaxin-like protein that shares only about 20% amino acid identities with vertebrate metaxins 1 and 2. But the vertebrate pattern of α-helical secondary structure is highly conserved in the plant and bacterial proteins (Adolph, 2020b). Alpha-helical regions, nine for plants and eight for bacteria, are the dominant feature. For protist metaxin-like proteins, the pattern of nine α-helices also resembles the pattern for human metaxin 1 (Adolph, 2021). The protists, eukaryotic organisms that are neither animals, plants, nor fungi, include animal-like protists such as *Amoeba proteus* (Amoebozoa) and plant-like protists such as the alga *Chlorella variabilis* (Chlorophyta). But even with this diversity, the secondary structures of all protist metaxin-like proteins are dominated by α-helices, with a pattern similar to other organisms.

Vertebrate metaxin 1, but not metaxin 2 or metaxin 3, was found to possess a transmembrane α-helical segment that largely coincides with the C-terminal helix H9. It was postulated that the helical segment anchors the protein to the outer mitochondrial membrane, where it functions in protein import into mitochondria. The transmembrane segment is not unique to vertebrate metaxin 1, and was found to be present in metaxin 1 homologs of invertebrates and the metaxin-like proteins of plants and protists, but not bacteria. The study reported here has revealed that MTXa proteins of fungi also have a transmembrane helix near the C-terminus, but MTXb proteins do not. This result was found for MTXa and MTXb proteins of fungi of different taxonomic divisions. Examples of Ascomycota fungi having MTXa proteins with a transmembrane helix near the C-terminus are the fission yeast *Schizosaccharomyces pombe* and *Aspergillus fumigatus*, a pathogenic fungus of humans. Basidiomycota mushrooms with the MTXa transmembrane helix include the cultivated white mushroom *Agaricus bisporus* and the wood rot mushroom *Heliocybe sulcata*. Further examples are *Allomyces macrogynus* and *Catenaria anguillulae* (Blastocladiomycota), *Rhizoclosmatium globosum* (Chytridiomycota), *Absidia repens* (Mucoromycota), as well as *Piptocephalis cylindrospora* and *Dimargaris crystalligena* (Zoopagomycota).

### 3.5. Relationship of Sam37 and FAXC Proteins to MTXa and MTXb Proteins

Experimental evidence has provided a detailed understanding of the role of Sam37, a protein of the outer mitochondrial membrane of yeast (*Saccharomyces cerevisiae*), in the import of β-barrel membrane proteins into yeast mitochondria. As shown in Figure 2, Sam37 possesses dominant Tom37 (E-value: 1.29e-39) and Tom37_C (4.85e-72) domains. These define the protein as Sam37. But Sam37 also has less dominant GST_N_Metaxin and GST_C_Metaxin domains, in common with the metaxins and metaxin-like proteins. The presence of GST_N_ and GST_C_Metaxin domains means that Sam37 is related to the metaxins, but is not in the same category as MTXa and MTXb proteins with GST_Metaxin domains as the major domains.

Sam37 is also an α-helical protein with a pattern of α-helices similar to the metaxins and metaxin-like proteins. The α-helical secondary structure supports the idea that Sam37 is a metaxin-related protein. Figure 5 includes a multiple sequence alignment of Sam37 with human metaxins 1 and 2, as well as MTXb of *Schizosaccharomyces pombe*. Sam37 is seen to possess an α-helical structure like that of the human metaxins and the fungal metaxin-like protein. Helices H1 through H8 are highly conserved in all of the amino acid sequences included in Figure 5. Human metaxin 2 has the additional N-terminal helix H*. MTXb of *Schizosaccharomyces pombe* also has an extra N-terminal helix. Human metaxin 1 has the additional C-terminal helix H9. Sam37 has a C-terminal helix that largely aligns with H9 of human metaxin 1.

Figure 6 shows amino acid sequence alignments of three metaxin-like proteins, MTXa, MTXb, and FAXC, found in the fungus *Neocallimastix californiae* of the Neocallimastigomycota division. Even though the three metaxin-like proteins share very little amino acid sequence homology (see section 3.2), the alignments demonstrate the high level of conservation of α-helical segments. The alignments display a similar pattern of helices as human metaxin 1 and human metaxin 2 included in the figure. In particular, helices H1 through H8 are present in all sequences.

Numerous other fungi also possess both MTX and FAXC proteins. Examples include *Piromyces finnis* (Neocallimastigomycota), found like *N*. *californiae* in the digestive tract of large herbivores, *Gonapodya prolifera* (Chytridiomycota), an aquatic fungus or water mold, *Heliocybe sulcata* (Basidiomycota), a mushroom that produces brown rot on wood, and *Powellomyces hirtus* (Chytridiomycota), a chytrid fungal species found in soil.

### 3.6. Phylogenetic Analysis of Fungal MTXa and MTXb Proteins

The phylogenetic tree in Figure 7 demonstrates that MTXa proteins and MTXb proteins are distinct categories of metaxin-like proteins. They are also separate categories compared to the human metaxins and Sam37, a yeast protein that shares features with the metaxins and metaxin-like proteins. Results like those in Figure 7 show that the vertebrate metaxins and SAM37 proteins are related by evolution to the metaxin-like MTXa and MTXb proteins of fungi. These analyses also suggest that the fungal metaxin-like proteins and vertebrate metaxins, as well as the Sam 37 proteins, evolved from a common ancestor.

The major protein domains of the Sam37 protein of *Saccharomyces cerevisiae* are the Tom37 and Tom37_C domains. GST_N_Metaxin and GST_C_Metaxin domains are also present, and therefore Sam37 has a domain structure related to that of the metaxins. Figure 7 includes *S. cerevisiae* Sam37, as well as the Sam37 proteins of other yeasts or fungi that are in the same family, Saccharomycetaceae, and are therefore closely related to *S. cerevisiae*. The Sam37 proteins form a separate group or cluster that is phylogenetically related to the MTXa, MTXb, and human metaxin proteins. Although the other yeast species are in the same family as *Saccharomyces cerevisiae*, there are some differences in their SAM37 proteins. For example, the *Candida*, *Debaryomyces*, and *Kluyveromyces* species have Sam37 proteins that lack the GST_C_Metaxin domain.

Figure 8 shows the phylogenetic relationships between three metaxin-like proteins MTXa, MTXb, and FAXC of *Neocallimastix californiae* and *Piromyces finnis*. These two species are members of the taxonomic division Neocallimastigomycota. Also included in the figure are the MTXa and FAXC proteins of *Anaeromyces robustus* (Neocallimastigomycota) and the MTXa and MTXb proteins of *Batrachochytrium dendrobatidis* (Chytridiomycota). The fungi of the Neocallimastigomycota division were previously part of the Chytridiomycota division, until that division was divided into Chytridiomycota, Neocallimastigomycota, and Blastocladiomycota divisions.

The phylogenetic analysis in Figure 8 shows three separate groups of metaxin-like proteins that represent the MTXa, MTXb, and FAXC categories. The *Neocallimastix* and *Piromyces* species in the figure have metaxin-like proteins in all three groups. These results demonstrate that the MTXa, MTXb, and FAXC proteins represent three phylogenetically distinct categories of proteins. This conclusion is supported by amino acid sequence alignments, as discussed in section 3.2, which show that there is little homology between MTXa and b and FAXC proteins of the same fungus. For example, *Neocallimastix californiae* MTXa shows only 16% amino acid identities when aligned with MTXb and 18% with FAXC, while MTXb aligned with FAXC shows only 14% identities.

Vertebrate metaxin 3 proteins are also included in the phylogenetic tree of Figure 8. Besides human metaxin 3, these consist of the metaxin 3 proteins of zebrafish (*Danio rerio*) and frog (*Xenopus laevis*). Human metaxin 3 shares 45% amino acid identities with human metaxin 1 and 22% with human metaxin 2. The vertebrate metaxin 3 proteins in Figure 8 indicate that the fungal MTXa and b and FAXC proteins are related by evolution to the vertebrate metaxins, and that the vertebrate metaxins and fungal proteins have descended from a common ancestor. These conclusions are also mentioned in the discussion above of Figure 7.

### 3.7. Genes Adjacent to the Multiple Metaxin-like Protein Genes of Fungi

In vertebrates, the genes that are next to the metaxin genes are highly conserved among different vertebrates. For human and mouse, the metaxin 1 genes are flanked by genes for thrombospondin 3, an extracellular matrix protein, and the lysosomal enzyme glucocerebrosidase. Human and mouse metaxin 2 genes are adjacent to a group of homeobox genes encoding transcription factors that regulate development. And the human and mouse metaxin 3 genes are between genes for thrombospondin 4 and cardiomyopathy associated 5 protein. Database searches reveal similar genomic regions for the metaxin 1, 2, and 3 genes of many other vertebrates.

With fungi, comparison of the genes immediately adjacent to the MTXa and MTXb genes of the same fungi shows that the MTXa and MTXb genes have different genomic contexts. And the neighboring genes in each case are different from the vertebrate genes. In addition, the genes that neighbor the MTXa and MTXb genes of different fungi are not well conserved. The exceptions are fungi that are taxonomically similar. For the example of *Schizosaccharomyces pombe*, the protein-coding genes adjacent to the MTXa gene on chromosome II are: rRNA processing protein Tsr2/recombination protein Saw1/phosphatidic acid phosphatase/**Metaxin-like Protein MTXa**/secretory component protein/translocation subcomplex subunit Sec66. This genomic region is very different from the *Schizosaccharomyces pombe* MTXb genomic region on chromosome I: protein mug74/mediator complex subunit Srb9/**Metaxin-like Protein MTXb**/pho88 family protein/WD repeat-containing protein Atg18.

Another example that demonstrates the differences in neighboring genes is *Talaromyces marneffei* (Ascomycota), an opportunistic fungal pathogen of humans. The genes adjacent to the MTXa gene are: ankyrin repeat-containing protein/β-D-glucoside glucohydrolase/**Metaxin-like Protein MTXa**/plasma membrane low affinity zinc ion transporter/metabolite transport protein. For the MTXb gene, the neighboring genes include: hypothetical protein/**Metaxin-like Protein MTXb**/copper-activated transcription factor GRISEA/DUF543 domain protein. The examples of *S. pombe* and *T. marneffei* show that different genes are adjacent to the MTXa and MTXb genes, and that the MTXa and MTXb genomic regions are not highly conserved between different fungi.

